# In vivo profiling of site-specific human cancer cell states in zebrafish

**DOI:** 10.1101/2021.06.09.447621

**Authors:** Dagan Segal, Hanieh Mazloom-Farsibaf, Bo-Jui Chang, Philippe Roudot, Divya Rajendran, Reto Fiolka, Mikako Warren, James F. Amatruda, Gaudenz Danuser

**Affiliations:** Department of Bioinformatics, UT Southwestern Medical Center, Dallas, TX; Department of Cell Biology, UT Southwestern Medical Center, Dallas, TX; Department of Pathology and Laboratory Medicine, Childrens Hospital Los Angeles, University of Southern California Keck School of Medicine, Los Angeles, CA; Cancer and Blood Disorders Institute, Childrens Hospital Los Angeles; Keck School of Medicine, Los Angeles, CA

## Abstract

Tissue microenvironments affect the functional states of cancer cells, but determining these influences *in vivo* has remained a significant challenge. We present a quantitative high-resolution imaging assay of cancer cell morphology in zebrafish xenografts to probe functional adaptation to variable cell extrinsic cues and molecular interventions. We focus on Ewing Sarcoma, a pediatric cancer driven by a single oncogenic fusion protein EWSR1-FLI1, and with little to no additional somatic mutations, making it a prototypical form of cancer whose adaptation to microenvironments is likely driven by acute, non-genomic mechanisms. Using computer vision analysis of 3D cell shapes, we find systematic shifts in the distribution of cell morphotypes between distinct sites in the fish embryo. We also find site-specific morphological responses to differential expression of EWSR1-FLI1. Combining these data we propose a model where Ewing Sarcoma cancer cell plasticity is sensitive both to expression fluctuation of EWSR1-FLI1 and signals from the surrounding tissue microenvironment, with either or both factors possibly contributing to the oncogenic potential of these cells.

## Introduction

Metastasis is the primary cause of cancer-related deaths [1]. The ability of metastatic cancer cells to arrive, survive and thrive in diverse tissue microenvironments (MEs) is a testament to their adaptability. Despite decades of research, our understanding of the mechanisms of metastasis has many remaining gaps, making this the single biggest obstacle to successful cancer treatment [1, 2]. Among the open questions on metastasis is the identification and systematic probing of the functional adaptation of cancer cells to specific tissue microenvironments.

Historically, cancer research has predominantly relied on the mouse for investigation of organotropic behaviors [3]. However, for the task of probing functional adaptation of cancer cells in different tissue environments, this model has several limitations. Cellular responses are dynamic and heterogeneous, and meaningful functional plasticity is super-imposed by cell-to-cell variation that is unrelated to organotropic processes. Statistical distinction of these sources of cellular heterogeneity require a large pool of single cell measurements in the intact cellular microenvironment, which is nearly impossible to accomplish in the mouse. Moreover, the mouse does not provide easy access for live cell imaging in most organs, limiting the diversity of microenvironments that can be studied. As an alternative, 2D and 3D cell culture have been exploited for studying cancer cell heterogeneity [4, 5], but these systems lack the relevant micro-environmental complexity of an organism for understanding functional adaptation.

Here, we propose human cancer cell xenografts in zebrafish larvae as a powerful compromise between the competing quests for environmental diversity, experimental access, and sufficient throughput to acquire meaningful descriptions of cancer cell adaptation [5–8]: (1) Zebrafish larvae contain various complex tissue microenvironments, with tissue mechanics, physiology, and disease susceptibility that is highly conserved with those of humans [6, 9]. (2) The optical transparency of zebrafish grants convenient experimental access to any metastatic site by high-resolution imaging, and (3) a single experimental session can yield data from numerous sites and xenografts, allowing statistically properly powered comparative studies.

This work sets the stage for development of human cancer xenografts in zebrafish as a model for unraveling mechanisms of cell adaptation during tumor progression and metastasis. As an initial readout of plastic cell behaviors, we focus on cell morphological variation. Cell morphology is the integrated response of cells to autonomous cues [10–15], mechanical and chemical interactions with the microenvironment [16–18], as well as a functional proxy for proliferation and cell death [19, 20]. Accordingly, pathologists have relied for decades on morphological patterns to stratify cancer disease states and to deliver specific diagnoses that guide treatment. With the recent development of machine learning, cell morphology is emerging as an increasingly quantitative correlate to particular molecular cell states [12, 21–25].

Studying cancer cell adaptation is of particular importance in pediatric cancers. While somatic variation, clonal heterogeneity and selection are thought to drive adaptation during metastasis of several adult cancers [26], many pediatric cancers exhibit a very low mutational burden beyond a single driving mutation [27, 28]. This suggests that non-genetic mechanisms may dominate the process of cell adaptation. In this work, we focus specifically on Ewing Sarcoma (EwS) as a pediatric cancer prototype. EwS is the second most common pediatric bone tumor, characterized by a chromosomal translocation fusing members of the *FUS/EWSR1/TAF15* family to *ETS* transcription factors, the most common of which creates the *EWSR1-FLI1* (*EF1*) fusion oncogene [29, 30]. As with most other pediatric sarcomas[31], presence of metastasis is by far the most prominent adverse prognostic factor in EwS [29]. The specific tissue environment appears to play a key role in the degree of malignancy in EwS [32–35], as well as other pediatric sarcomas [36, 37]. For example, pelvic primary tumors appear to correlate with poorer prognosis and are more prone to metastasis compared to other primary sites, and patients who have metastasis only to lung have improved survival compared to those with metastasis to bone or bone marrow [32].

Thus, we assembled a quantitative high-resolution imaging assay of EwS cell morphology in xenografts to probe the morphological variation as a proxy to functional adaptation in diverse microenvironments and experimental molecular interventions. Applying in house-written computer vision pipelines to 3D cell shape analysis, we found systematic shifts in the distribution of cell morphological states between seeding sites in the fish embryo, as well as between control expression and knockdown of *EWSR1-FLI1* in an environmentally sensitive fashion. We also found that proliferative but not apoptotic states of EwS cells differed between the tissue microenvironments. We propose a model where EwS cancer cell plasticity is sensitive both to fluctuations of EWSR1-FLI1 and signals from the surrounding tissue microenvironment, with either or both of these factors possibly contributing to metastatic potential.

## Design

### 3D shape analysis captures functional heterogeneity in Ewing Sarcoma cells

We present an experimental and analytical pipeline to identify single-cell morphologies within tissues of zebrafish larvae and identify differences in morphological cell states (Figure 1). By analysis of geometric features extracted from high-resolution 3D image volumes, we examined morphological heterogeneity as a function of cell type, oncogene expression level, and tissue microenvironment. To allow single cell segmentation and shape analysis, mixtures of cells tagged with either a red-emitting (90-99% of cell mixture), or a green-emitting filamentous actin marker F-Tractin (1-10% of cell mixture) were injected into 2-day old zebrafish larvae at one of three seeding sites: the hindbrain ventricle (HBV), perivitelline space (PVS), and, by rapid passive transport during injection, the caudal hematopoietic tissue (CHT). This allowed us to investigate morphological and functional adaptation of cell population to distinct microenvironments. For example, the HBV are cavities which contain cerebrospinal fluid, rich in hormones, proteoglycans, and ions [38–40], the PVS is a sandwiched, avascular region between two malleable membranes surrounding the yolk [41] and the CHT is a compact, thin, crowded region containing multiple cell types, including endothelial cells, stromal cell and hematopoietic cells for which it serves as a stem cell niche [42–44].

**Figure 1:**
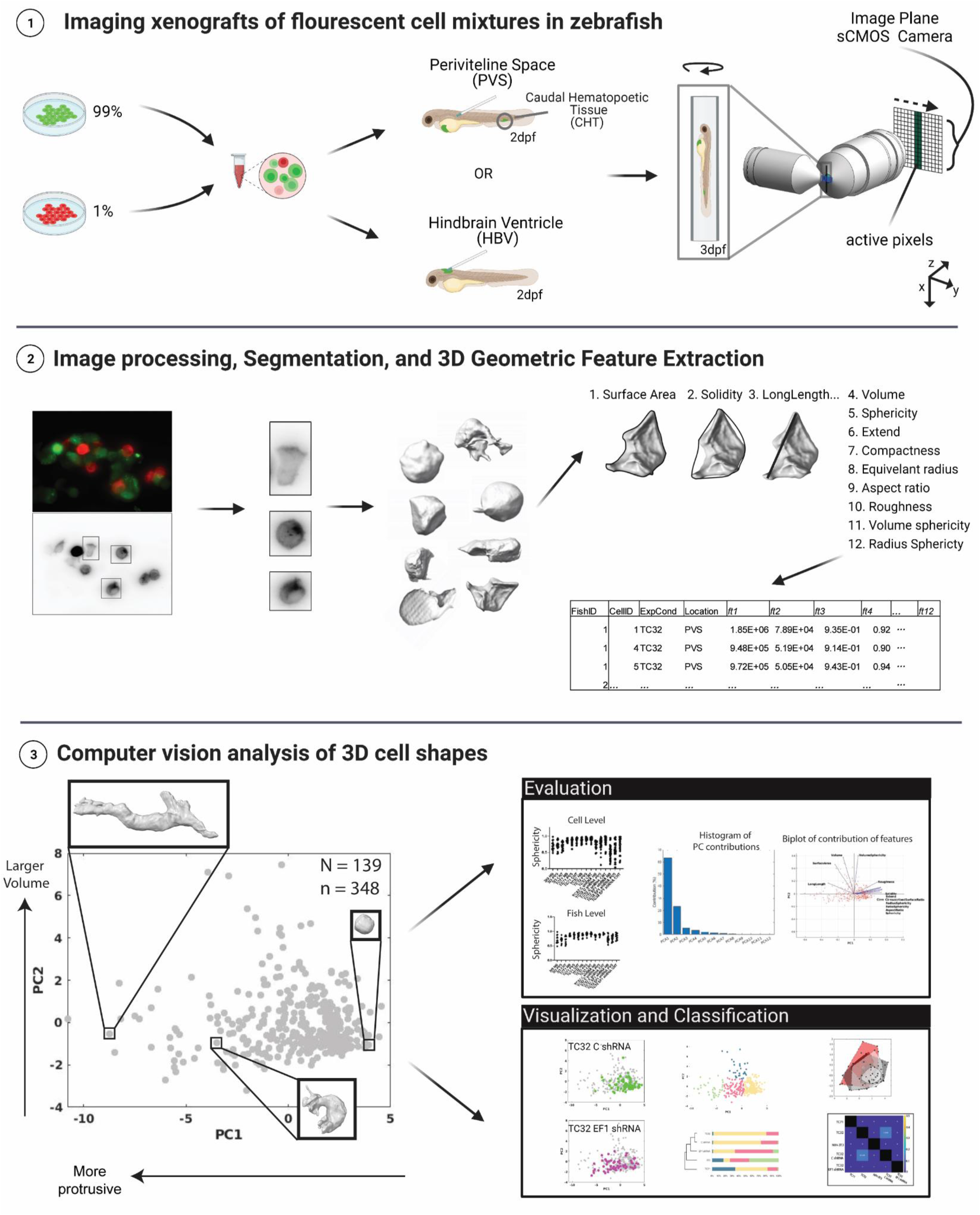
3D cell shape analysis in cancer cell xenografts in zebrafish larvae. We present an experimental and analytical pipeline to identify single-cell morphologies within tissues of zebrafish larvae and identify differences in morphological cell states by computer-vision analysis of 3D cell shape, as follows. (1) In order to distinguish shapes of individual cells, we prepared a mosaic cell mixture consisting of 90-99% of cells labeled with a green-emitting fluorophore and 1-10% of cells labeled with a red-emitting fluorophore, or vice versa. We then injected the cell mixture into the Perivitelline space (PVS) or Hindbrain Ventricle (HBV) of 2-day-old zebrafish larvae. For PVS injections, we selected specifically for injected larvae which also have xenografted cells in the caudal hematopoietic tissue (CHT) immediately following injection. Fish larvae were fixed at 3 days post fertilization (dpf) and imaged using an Axially Swept Light Sheet Microscope (ASLM), which is optimized for maximum throughput, resolution, and flexibility of sample orientation. High axial resolution is obtained by combining a thin, moving light-sheet with a synchronized rolling shutter (active pixels) that blocks out of focus fluorescence on the image plane on an sCMOS Camera. (2) Individual cells were computationally extracted from the image data of sparsely labeled cells, which after image processing are segmented into 3D volumes. We then measured 12 global 3D geometric features as listed, and each cell was indexed according to fish, location, and cell type. (3) Using Principal Component Analysis (PCA) we captured the heterogeneity of morphological cell states (N=number of fish, n=number of cells). PC1 captured the protrusive state of the cell, while PC2 was influenced largely by volume (insets). We evaluated our analytical pipeline and used several visualization and classification techniques to capture differences among heterogeneous single-cell data.

24 hours following injection, chemically fixed zebrafish larvae were mounted in 1% low-melting agarose in Fluorinated Ethylene propylene (FEP) tubes, which allows for rotation of the sample in order to optimize the imaging conditions at the sites of interest. Images within the three seeding sites were acquired using an Axially Swept Light-sheet Microscope (ASLM) [45, 46], which is a high-resolution modality of light-sheet fluorescence microscopy (LSFM) that offers near-isotropic resolution over large sampling volumes. Isotropic resolution is essential for an unbiased analysis of 3D cell geometry [12]. For the presented study we imaged ~190 fish larvae. After semi-automated image processing to identify image sub-volumes containing cells suitable for shape analysis (see Methods), we extracted the volumes of ~640 individual cells using a tailored version of an automated segmentation pipeline developed previously [12]. Each computationally defined surface was visually inspected to eliminate segmentation artifacts. The final dataset encompassed ~350 validated cell volumes from ~140 fish. For each volume, we measured 12 geometric features (Figure 1, Table S1). To test for batch effects, we calculated the median values of each feature per fish (Figure S1a). While the geometric features showed substantial variance, even within bins that distinguish cell type, oncogene expression and seeding site, the variance of the features among fish was decreased, suggesting that batch effects across larvae are modest. Significant shifts between bins, even for a single feature at the fish level, indicate that cell type, molecular condition, and seeding site play a role in the control of cell morphogenesis.

To capture differences in the morphology states of bins we switched to multivariate analysis of the complete feature sets. Pairwise comparisons of individual features such as sphericity and volume could not distinguish many cell types or molecular states (Figure S1b). To visualize and reduce the dimensionality of the 12-dimensional feature space we applied principal component analysis (PCA) (Figure 1). The first two principal components (PCs) captured 86% of the heterogeneity within the data, indicating a high-level potential for dimensionality reduction of the feature space (Figure S1c). Indeed, PC1 and PC2 together captured morphological differences between all cell types and molecular conditions (Figure S1d). As a negative control we also compared WT cells to cells of the same type expressing scrambled shRNA vectors and found no difference between the conditions. We wondered which geometric properties were most important for a EwS cell morphology analysis. To address this, we projected the unit vectors of the twelve individual features into the (PC1, PC2)-space. PC1 was dominated by Sphericity and Aspect Ratio, thus capturing the protrusive state of the cell. PC2 was dominated by Volume and Surface Area, thus capturing the size and rectangularity of a cell (Figure S1e).

Together, these outcomes led us to conclude that a 2D linear projection of the original feature space would be sufficient for the analysis of meaningful morphological differences between EwS cells.

## Results

### EwS cell morphologies are different from fibroblasts and differ between cell lines

Equipped with the described pipeline, we examined single cell morphologies of two patient-derived Ewing Sarcoma cell lines, TC32 and TC71, *in situ* in zebrafish larvae (Figure 2a). We further included analysis of NIH-3T3 fibroblasts as a reference of non-transformed cells. Previous work in 2D cell culture reported strong morphological changes in response to knock-down of EF1 gene expression, which were also accompanied by shifts in proliferation but not in cell survival [47]. Therefore, we wondered whether EF1 expression changes would trigger similar responses in our *in situ* 3D imaging system. To accomplish this, we injected TC32 Ewing sarcoma cells expressing a doxycycline-inducible scrambled (C shRNA) or EF1 (EF1 shRNA) targeting shRNA and cytoplasmic GFP tag and with (1-10%) or without (90-99%) ubiquitously driven F-Tractin-MRuby2. Visual inspection of 3D renderings of segmented cells already showed distinct differences between NIH-3T3 and EwS cells, and between cells with wild-type or knockdown levels of EF1 expression, pooled from all three tissue MEs (Figure 2b). Projection of the data into the (PC1,PC2) space confirmed the visual impression of strong morphological shifts between these cell types and conditions (Figure 2c,d).

**Figure 2:**
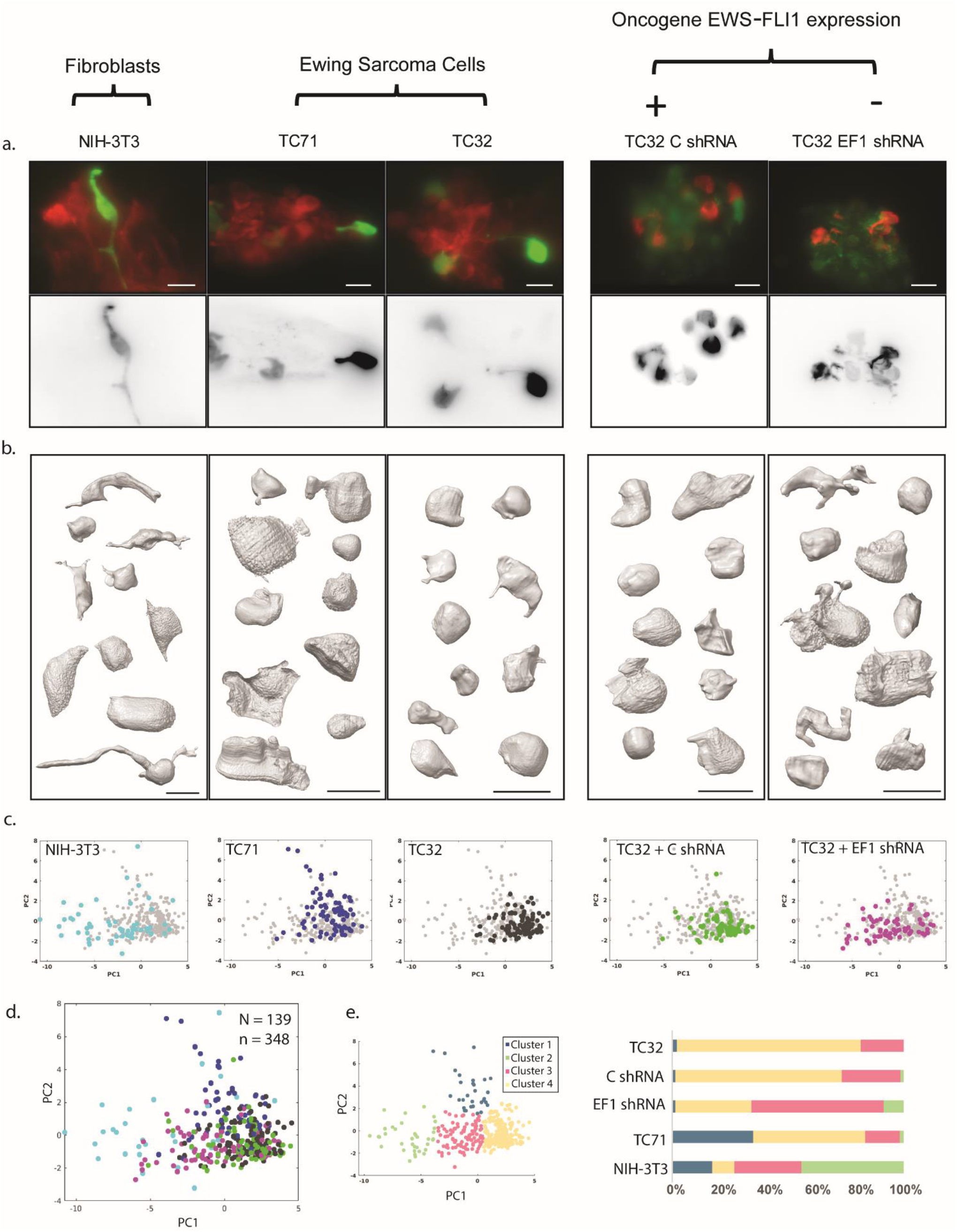
Classification of morphologies captures heterogeneity of cell states in EwS cells. (a, left) EwS (a, TC71 and TC32) or fibroblast (a, NIH-3T3) cells expressing FTractin-GFP (green in upper panel, grey in lower panel) or FTractin-mRuby2 (red), in zebrafish HBV (3dpf). (a, right) TC32 cells expressing a doxycycline-induced cassette which includes a cytoplasmic GFP (upper panel, green) as well as (scrambled shRNA (C shRNA) or shRNA targeting EF1 (EF1 shRNA). A subset of cells also express FTractin-mRuby2 (red in upper panel, grey in lower panel), allowing for sparse labeling of single cell morphologies. (b) 10 representative 3D cell renderings are displayed within each experimental group, pooled from all three tissue MEs. Note the distinction in cell shape and size particularly in NIH-3T3 (left-most panel) and TC32 EF1 shRNA (right-most panel) compared to other experimental groups. (c-d) PCA of all experimental groups shown individually (c) or together (d). (e) K-means based clustering allows visualization of relationships between experimental groups. Note distinctions in all experimental groups. Scale bars 20μm.

To better reflect the variation of cell shapes, we visualized differences between bins by assigning cells to one of four characteristic clusters in (PC1,PC2) space. We identified the clusters by K-means and determined the number k of clusters subject to maximizing the silhouette score in the range [3, 10] clusters (Figure 2e). Clusters 4, 3, and 2 break the data set primarily along the PC1 axis, representing increasingly more protrusive cell shapes, respectively. Cluster 1 is characterized by a shift in PC2, representing larger, more rectangular cells. Comparison of the cluster assignments demonstrates stark differences between cell types. Fibroblasts are heterogeneous, with sampling of all four clusters. In contrast, TC71 cells dominate Cluster 1, while TC32 cells predominantly occupy Clusters 3 and 4. TC32 cells with a knock-down of EF1 expression displayed an increase in occupancy of Clusters 2 and 3, at the expense of Cluster 4 occupancy, i.e. lower EF1 expression overall yields more protrusive cells. This observation is consistent with 2D cell culture studies describing the cells with EF1 knock-down as more ‘mesenchymal’ [47]. Of note, exclusion of NIH-3T3 as an outlier group resulted in only a minor shift in PCA space and relative contribution of features (Figure S1f), suggesting that the PCA of the 12 global 3D features constitutes a robust space for the definition of cell morphological classes and is not dominated by the fibroblast distribution. Overall, these results demonstrate the applicability of 3D cell shape analysis in capturing differences across cell types and experimental conditions, and demonstrate that EF1 oncogene levels affect EwS cell morphology *in vivo*.

### Tissue microenvironment affects EwS cell morphology and function

In order to examine whether EwS cells functionally adapt to the tissue ME, we first compared proliferation and cell death states between HBV, PVS, and CHT (Figure 3a) by immunostaining for the proliferative marker Phospho-Histone 3 (PH3) or the apoptotic marker TUNEL (Figure 3b-c), or by measuring change in cluster size over 4 days (S2a-c). While apoptosis was similarly low across all three tissue MEs (less than 10% on average, Figure 3d), proliferation and growth rates were higher in PVS compared to the other sites (Figure 3e, Figure S2d). Note that the number of cells greatly varied between regions (Figure S2e), a difference that likely arose from differences in initial seeding rather than cell growth. However, no clear correlation was seen between proliferation rate and cluster size (Figure S2f) or fold growth and initial cluster size (Figure S2a-c), suggesting cluster size may not be a major driver of differences in cell growth.

**Figure 3:**
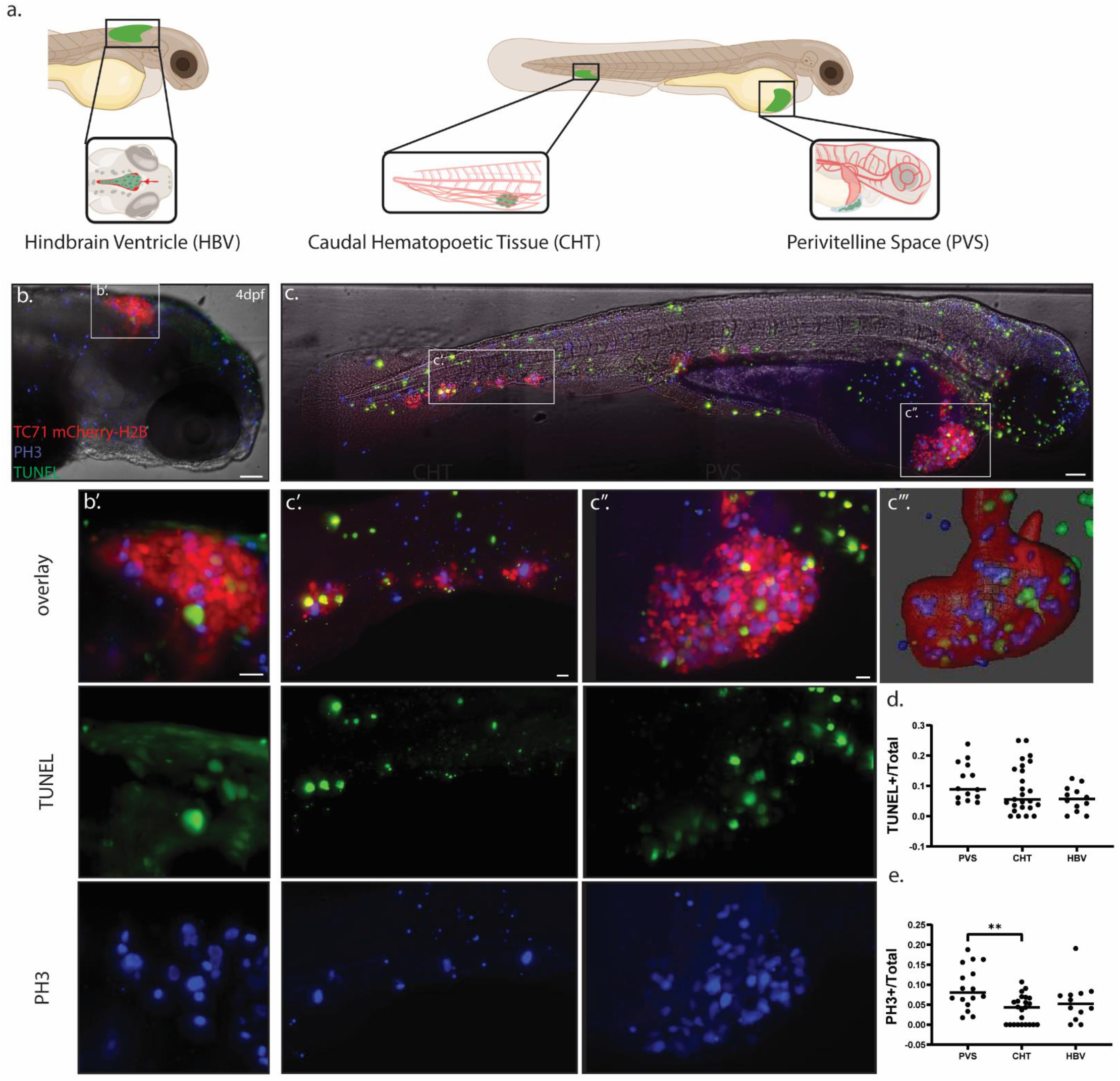
Proliferation but not apoptosis change as a function of tissue microenvironment. (a) Comparison of single cell morphologies were made as a function of seeding site within the zebrafish larvae, each representing a distinct tissue ME. The hindbrain ventricle (HBV) is a mostly fluid-filled compartment (a), the perivitelline space is a small, compressed region sandwiched between two membranes surrounding the yolk, and the caudal hematopoietic tissue (CHT) is a narrow region rich in vasculature. **(b-c)** TC71 cells expressing nuclear marker mCherry-H2B (red), within the HBV (inset, enlarged in b’), CHT (inset, enlarged in c’), or PVS (inset, enlarged in c’’), of fish larvae (4dpf). Fish are immunostained for proliferation marker PH3 (blue) and apoptotic staining TUNEL (green). Measurements are made on 3D volumes, as rendered in c’’’. **(d-e)** Apoptotic rate is similarly low across all three regions (d), while proliferation rate appears to be highest in PVS compared to other regions (e). Scale bars 100 μm (b,c) and 20 μm (b’-c, c”).

We next examined single cell morphological state variations between distinct tissue MEs (Figure 4). Bagplots (i.e. bivariate boxplots) [48] enabled visualization of differences in bin distributions in the PC space. Differences were further quantified by assembly of similarity matrices based on pairwise permutation tests between regions. Examination of NIH-3T3 fibroblasts revealed highly heterogeneous cell morphologies, with largely overlapping distributions in all three seeding sites, suggesting that these non-transformed cells lack sensitivity to environmental cues (Figure 4a). In contrast, the EwS cell lines TC32 (Figure 4b) and TC71 (Figure 4c) showed some distinction in the distributions of cell morphologies as a function of seeding site. For example, TC32 cells in the PVS and CHT were largely overlapping, but cells in the HBV shifted to the upper left in PC space, representing more protrusive, larger cells. TC71 cell distributions in the PVS and HBV shifted left in PC space compared to CHT (Figure 4c), suggesting that these cells stayed more rounded in the CHT. Importantly, the two EwS cell lines displayed differential adaptation patterns to these seeding sites indicating that their oncogenic transformations yield distinct tissue-specific specialization programs.

**Figure 4:**
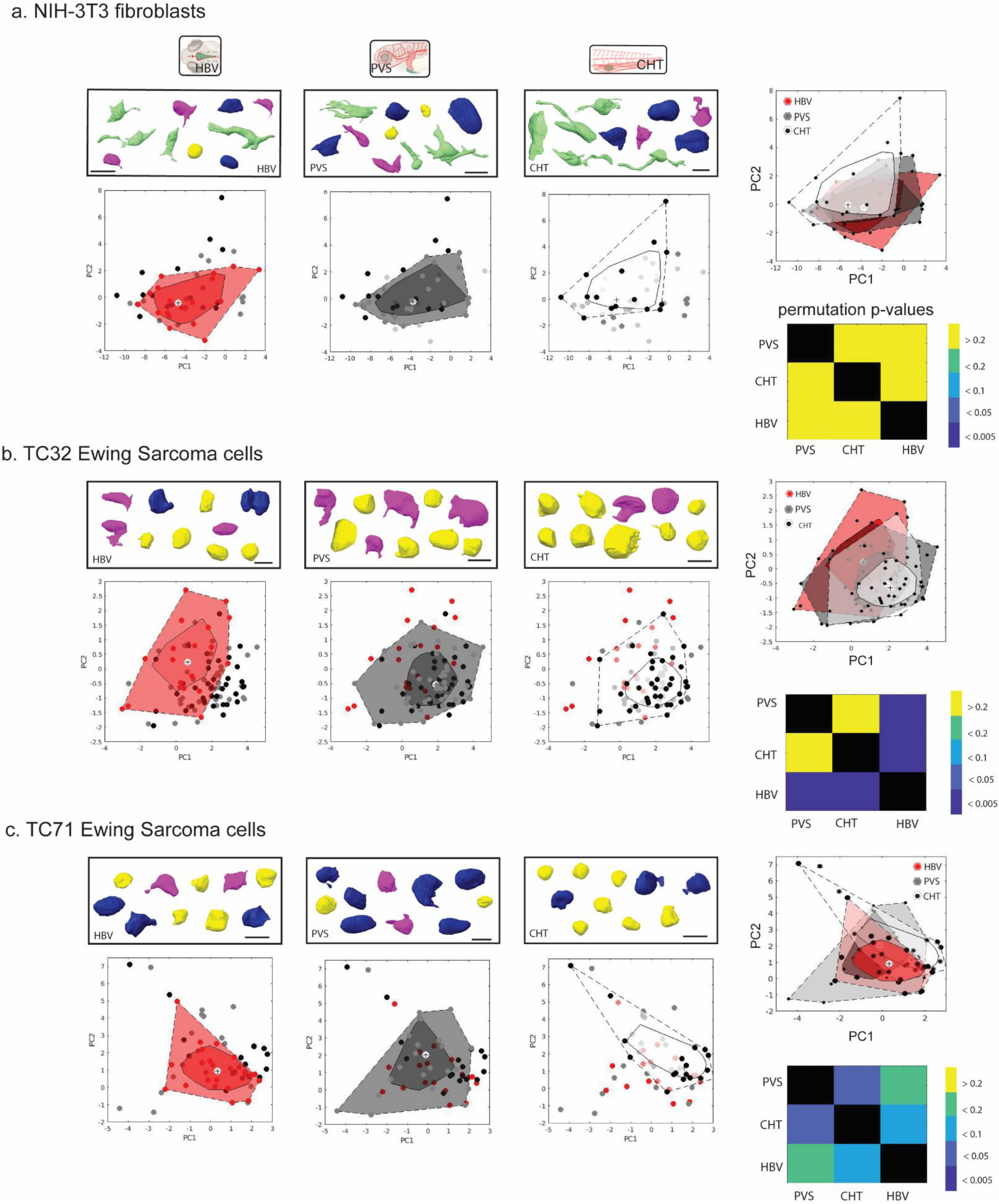
Tissue microenvironment affects EwS cell morphologies. (a-c) 10 representative cell renderings of EwS cell lines TC32 and TC71 and fibroblast cell line NIH-3T3 are shown, color-coded by k-means-based cluster from Figure 2e (black frames, upper part of panels). Bagplots below display PCA of each cell line as a function of region (HBV-red, PVS, black, CHT, white). The inner polygon is constructed on the basis of Tukey depth, and contains at most 50% of the data points. The outer circle encompasses all data excluding outliers. White plus sign represents the Tukey depth median. Lower right panel shows a similarity matrix based on pairwise permutation tests. Note that relative to NIH-3T3 which shows high similarity in morphological cell state among all three regions (a), TC32 (b) and TC71 (c) show some tissue-dependent specialization of cells. Scale bars: 20 μm.

Common to both TC32 and TC71 cell lines is the tendency to become more protrusive in the HBV. This was unexpected, given that the fluid-filled HBV compartment is likely softer than PVS and CHT, and thus is a mechanical environment usually associated with rounded cells [18]. On the other hand, this environment may be enriched in signaling cues that trigger neural differentiation [38, 39]. Closer examination of the architecture of this region revealed that EwS cell clusters within the HBV organized in pseudorosette-like spheroids, with protrusions pointing inwards and cell bodies occupying the cluster periphery (Figure 5a-c, Movie S1). Notably, pseudorosettes have been described clinically in a subset of Ewing sarcomas classified previously as peripheral Primitive Neuroectordermal tumors (ppNETs) [49–51], so named by pathologists according to their apparent neural-like differentiation [52]. Indeed, direct apposition of H&E stains of a human EwS tumor in soft tissue (Figure 5d) and TC71 cells expressing the fluorescent marker Cherry-H2B (Figure 5e) showed a striking similarity in the circular arrangement of nuclei associated with the formation of pseudorosettes. As expected, based on the cell morphological classification, these arrangements could not be observed with TC71 nuclei in the PVS or CHT (Figure S3a). In sum, these analyses demonstrate that a morphological classifier is a valid step towards the discovery of cell functional adaptation to MEs. They also show remarkable parallels between adaptive processes in human tissue and zebrafish, further validating the zebrafish model as a tool for studying organotropism of cancer cells.

**Figure 5:**
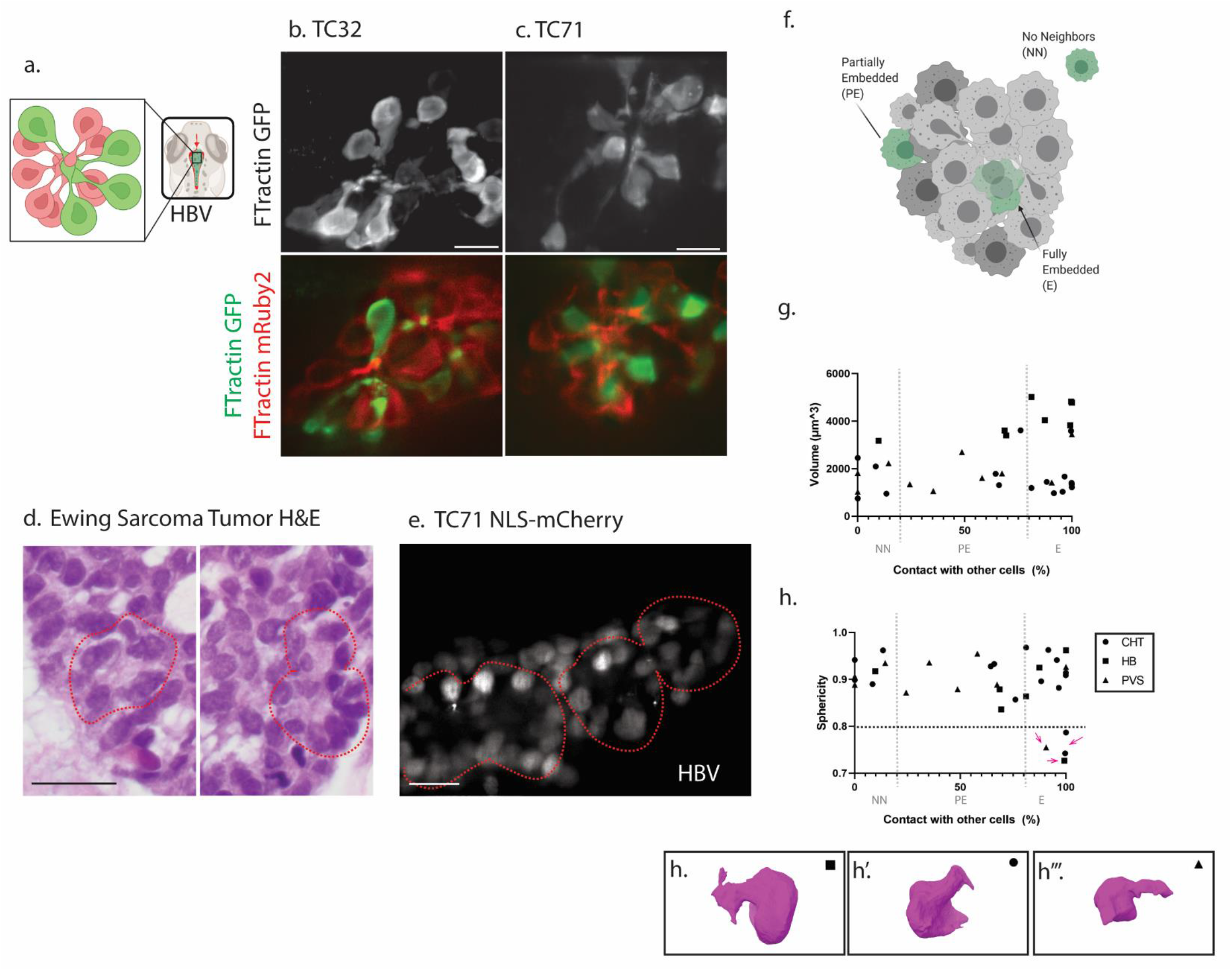
Tissue architecture affects cell state. (a) schematic of mixtures of flourescently labeled cells displaying pseudo-rosette-like formations formed by EwS cells specifically in hindbrain ventricles (HBVs). (b-c) Maximum intensity projections (upper panel) or single optical slices (lower panel) of TC32 and TC71 cells expressing Ftractin-GFP (grey, green) or F-Tractin-mRuby2 (red), in zebrafish larval hindbrain ventricle (3dpf). Note the pseudorosette-like polarization of cells, with cellular protrusions pointing towards a center of a sphere and cell bodies organized outwards. (d) An H&E stained histological slice from a human Ewing Sarcoma tissue sample from the soft tissue (sacral region) of a 20yo female, evaluated by a pathologist. Psuedorosettes structures outlined with dashed red line. (e) Single optical slice of TC71 cells expressing nuclear marker H2B-Cherry form ring-like structures (outlined with dashed red line) in the HBV which resemble those observed in some human Ewing Sarcoma tumors. (f) Cells within a cell cluster of like-cells are classified and analyzed according to percent contact with other cells, as no neighbors (NN, 0-20% contact with other cells), partially exposed (PE, 21%-81% contact), or fully embedded (E, 81-100% contact). (g-h) Proportion of contact with other TC32 cells is plotted against two dominant shape features, Volume (g) and Sphericity (h). While no clear relationship can be uncoupled from tissue ME for volume, a few embedded cells but not other groups show cells with distinctly low sphericity (>0.8, dashed line). Arrows indicate cells from all 3 regions with distinctive cell morphological states, rendered in h’-h’’’ and color-coded by k-means clustering in Figure 2e. Scale bars 20um.

Cell-cell interaction, cell-crowding, and position within a cell cluster or tissue all may affect cell function and thus the morphological state [53, 54]. We therefore tested whether the morphological state of EwS cells was affected by its position within a cell cluster. We distinguished cluster-peripheral and cluster-interior cells based on the fraction of the cell surface in contact with other cells in the cluster. For example, a cell may have few or no neighbors (NN,0%-20% of cell surface in contact with the cluster), be partially embedded within a cluster (PE,21%-80%) with one portion exposed to the tissue ME, or almost fully or entirely embedded within a cluster ((E,81%-100%), Figure 5f). We compared the degree of contact of EwS cells with other cells to two dominant shape features, sphericity and volume. At large, volume and sphericity are independent of the contact surface fraction. No correlation could be uncoupled from tissue ME for volume (Figure 5g,h; Figure S3b,c). However, the few cells with distinctly low sphericity were all fully embedded within the cluster, regardless of seeding site (Figure 5h, Figure S3c). Taken together, these results demonstrate a sensitivity of EwS cell morphological state to cell-extrinsic cues, demonstrating their potential for context-driven cell adaptation.

### Loss of EWSR1-FLI1 causes site-specific changes in cell morphology

Previous works have suggested that variation in EWSR1-FLI1 expression could drive differential cell function [55], including morphology[47]. However, these studies have so far been limited to 2D culture systems. Thus, we wondered how EWSR1-FLI1 expression shifts affect cell morphology, and whether putative morphological adaptation varies between the MEs presented by the three seeding sites. Using a dox-inducible EF1 knockdown system, we observed efficient partial reduction in EF1 (Figure S4a). However, associated changes in cytoskeletal organization were mild with no notable effects on cell shape (Figure S4b). This is in contrast to the significant shifts in cell morphology in the xenograft assays (Fig. 2).

We next examined whether loss of EF1 affected the cell morphological states in a tissue-sensitive manner. As with the wildtype version, TC32 cells expressing control (C) shRNA showed a distinct morphological shift of HBV relative to PVS and CHT, with cells shifting upward in PC2 ((Figure 6a-b, Figure S4c). The TC32 C shRNA cells showed additional distinction between PVS and CHT, with cells in PVS shifting to a relatively more protrusive state (left in PC1). Cells with an EF1 knock-down (EF1 shRNA) exhibited overall larger morphological heterogeneity and a significant shift of cells in all three regions towards a more protrusive state (Figure 6c-d, Figure S4c). Differences observed in the morphological states of (C) shRNA expressing cells between PVS and CHT became relatively blurred in EF1 shRNA group, while HBV cells remained distinctly higher in PC2 and volume. Pseudorosette-like structures formed also in HBV with EF1 low cells, despite altered morphology (Figure 6e, Movie S2), suggesting maintained sensitivity to the environmental cues of this environment. Examination of sphericity as a function of position within a cluster showed that upon EF1 knockdown cells with distinctly low sphericity could now also be observed with only partial contact to other cells (Figure 6f). This is in contrast to the requirement for a high degree of contact for TC32 and TC71 cells with a protrusive phenotype (Figure 5h, Figure S4c), suggesting that with decreased EF1 expression EwS cells become less sensitive to interactions with neighboring cells in the cluster. We conclude that reduced EF1 expression leads to an indiscrimately more protrusive morphological cell state with loss of sensitivity to some but not all environmental cues.

**Figure 6:**
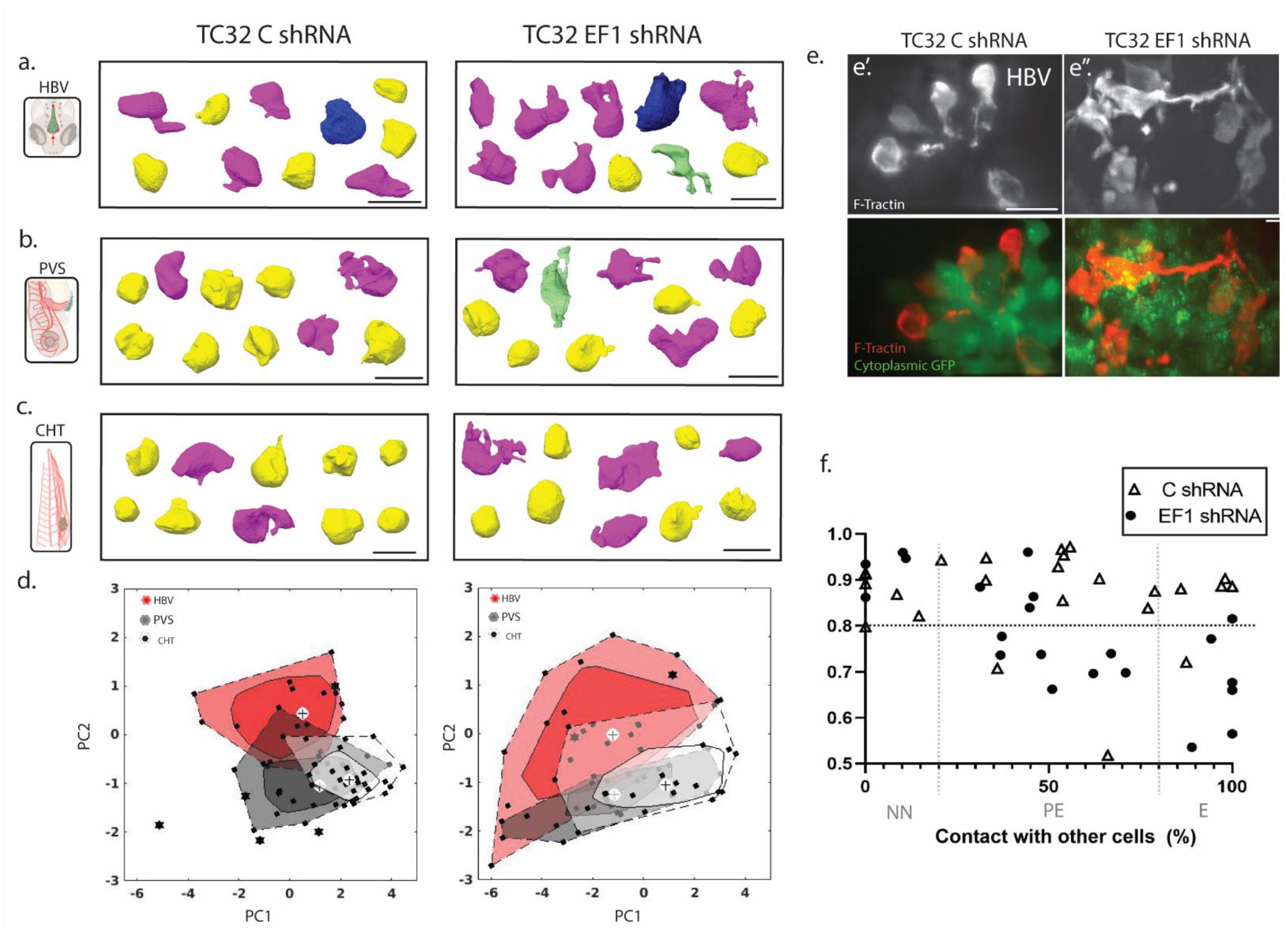
Knockdown of EF1 causes variable changes in morphologies as a function of tissue microenvironment. (a-c) Morphological cell states were examined as a function of tissue microenvironment (HBV, PVS, or CHT) in EwS cells expressing a dox-inducible scrambled (C, left) or EWSR1-FLI1 (EF1, right) targeting shRNA. Representative cell renderings from each group or color-coded by k-means clustering in Figure 2e. (d) Bagplots below display PCA of each cell line as a function of region (HBV-red, PVS, black, CHT, white). The inner polygon is constructed on the basis of Tukey depth, and contains at most 50% of the data points. The outer circle encompasses all data excluding outliers, which are marked by black asteriks. White plus sign represents the Tukey depth median. In C shRNA expressing cells, all 3 regions show distinct cell states in PCA space (left). Upon knockdown of EF1, cells are shifted to a more protrusive state and differences as a function of region are relatively reduced (right). (e) A subset of EwS C (left) or EF1 shRNA (right) expressing cells express F-Tractin mRuby2 (white in top panel, red in bottom panel) in addition to dox-inducible cytoplasmic GFP (green, bottom panel), allowing for labelling of individual cells. EwS cells in HBV can maintain a pseudorosette-like organization upon EF1 knockdown, despite altered cell morphologies. (see also movie S2). (f) Proportion of contact with other TC32 cells expressing C shRNA or EF1 shRNA is plotted against Sphericity. Upon Ef1 knockdown, cells with distinctly low sphericity (>0.8, dashed line) could now also be observed with only partial contact to other cells. Scale bar 20 μm.

## Discussion

In this study, we present a quantitative 3D imaging pipeline for the analysis of cell morphological adaptation to distinct microenvironments in zebrafish. Applied to human Ewing Sarcoma (EwS) xenografts we demonstrate that the pipeline captures systematic shifts in EwS cell morphological states as a function of cancer cell type, oncogene expression level, and location within the fish. We do not observe site-specific adaptation for xenografts composed of human fibroblasts, suggesting that organotropic behavior is an acquired feature during transformation.

Demonstrating *in situ* the plasticity of cancer cells in response to mechanical and chemical parameters has been difficult. Environmentally triggered variation must be gauged against a slew of stochastic cell autonomous cues that drive heterogeneous cellular responses. Our approach begins to address this very challenge by leveraging the combined advantages of the zebrafish model [6–9] and innovations in high-resolution imaging and computational image analysis, allowing for the throughput and resolution needed to uncover meaningful heterogeneity among noise. First, isotropic high-resolution image acquisition using ASLM enabled the segmenation and morphological characterization of cells in 3D with a sensitivity not achievable with other light-sheet imaging modalities, including the lattice light-sheet microscope [56, 57] (Figure S5a). Simulations of imagery with 3-fold reduced axial resolution yielded segmentation errors and marked shifts in PCA space, demonstrating the importance of high, near-isotropic resolution in 3D measurements (Figure S5b-c). Second, because of intrinsic cell-to-cell variation, the analysis has relied on a fairly large dataset. For example, random removal of 50% of the data caused loss of sensitivity to differences between experimental groups (Figure S5d). Third, the automation of the single-cell segmentation and shape feature definition using constant hyper-parameters across all image data sets guaranteed an unbiased analysis. Notably, detected differences between groups were overall robust within a 30% margin of controlparameter variation (Table S2). Overall, these technical developments provide a powerful basis for to study cancer cell adaptation.

We applied the pipeline to investigate evidence of extrinsically driven variation of EwS cell states, a form of cancer whose progression we hypothesize is strongly dependent on cell adaptation to the microenvironment. Indeed, we found that both environmental conditions of the host tissue as well as the location within the sarcoma cell cluster systematically affect EwS cell morphology (Figure 7). In contrast, non-transformed fibroblasts, while even more heterogeneous in morphology than EwS cells, showed no sensitivity for the tissue ME, suggesting that specialization for certain environments is a cellular feature acquired with the oncogenic transformation of EwS. Our results also show that expression levels of EF1 can regulate morphological cell state *in vivo*. We find that knock-down of the EF1 fusion protein leads to an overall shift towards more protrusive cell morphologies. This is consistent with previously reported observations of EwS in cell culture, which reported shifts in transcriptional and morphological cell states characteristic of epithelial-to-mesenchymal transitions (EMT) [47] and decreased proliferation [58, 59]. Perhaps more important than a global shift in cell morphology, our data show a homogenization of cell morphologies between tissue MEs and cluster locations upon EF1 knock-down.

**Figure 7:**
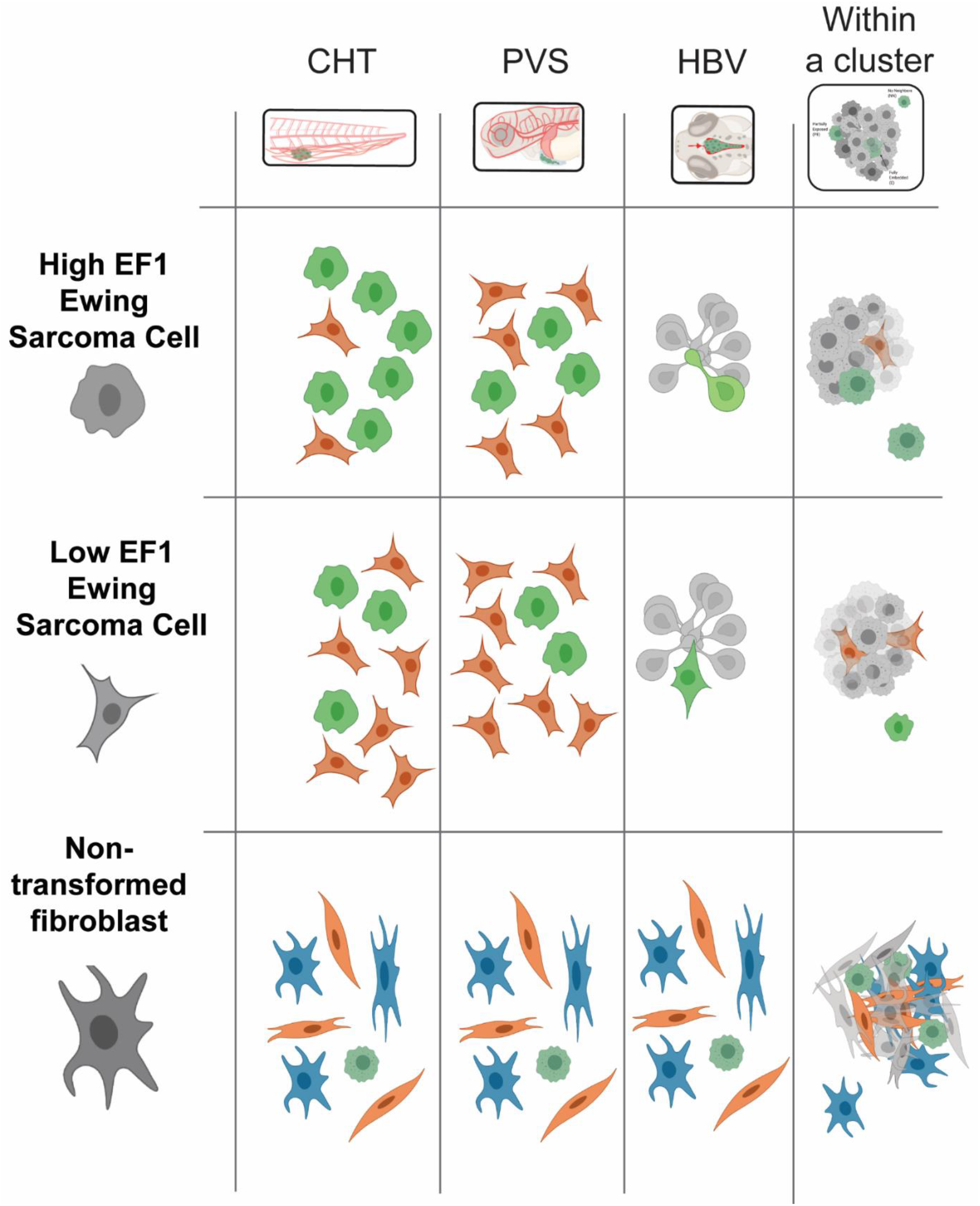
Ewing Sarcoma morphological cell state displays context-specific cellular adaptation. A schematic representing findings. High EF1 expressing EwS cells show a change in morphological state or organization according to seeding site or location within a cluster. Low expression of EF1 causes a shift towards more protrusive cell state and reduced sensitivity to some tissue environmental cues but not others, suggesting EF1 enables site-specific specialization of cancer cells. In contrast, non-transformed fibroblasts display highly heterogeneous cell morphologies regardless of extrinsic cues.

Based on these results, we propose EwS cells through expression of EF1 acquire a “high-plasticity cell state” (HPCS), as recently identified in lung cancer [60]. A transition into an HPCS allows EwS cells to adapt to distinct tissue microenvironments. Low expression of EF1 confers a loss of plasticity and reduced proliferation, which are properties associated with a more differentiated cell state. Our interpretation of EF1 as a plasticity factor is complementary to models in the EwS field, which suggests that modulation of levels of EF1 can enable switch-like behavior from proliferative to migratory states [47, 55]. It is the merit of the proposed zebrafish system offering different MEs rather than the monotonic conditions of *in vitro* cell culture that begins to unravel this potentially critical function of EF1 for disease progression. Future work will now systematically exploit this experimental system to determine the molecular underpinnings of an EF1-driven cell plasticity.

### Limitations and future prospects

While this work represents a significant advance in characterizing cell adaptation to tissue microenvironments, several limitations of the current approach must be considered. First, while morphology serves as a powerful functional output for cell state, it is a conglomerate of many mechanical [18, 61, 62], biochemical [13, 14], and stochastic [63] processes and therefore cannot inform on the specific pathways regulating cell behaviors such as plasticity. For a deeper understanding of the observed tissue-dependent cell differences we will now have to introduce molecularly-specific probes. Our pipeline already fulfills the requirements in terms of microscope resolution and sensitivity to perform analyses of subcellular molecular behaviors. Second, the throughput of the assay could be greatly enhanced if mosaic labeling was expanded to multicolor techniques such as brainbow [64]. Third, while the zebrafish embryonic xenograft model confers many advantages, it is limited to the short timescale of 2-10 days post-engraftment. To investigate longer term cell adaptations, complementary similar approaches should be developed for adult immunocompromised zebrafish [65], organoid [66, 67], and murine cancer models [68]. Finally, the potential of this system will have to be expanded for other cancer types.

Together the presented work establishes a framework for the statistically robust study of cancer cell plasticity in diverse microenvironmental cues, and led to the proposal of the EF1 fusion protein driving EwS disease as a plasticity factor.

## Methods

### Zebrafish husbandry

*Danio rerio* were maintained in an Aquaneering aquatics facility according to industry standards. Vertebrate animal work was carried out by protocols approved by the institutional animal care and use committee at UT Southwestern, an AAALAC accredited institution. WIK wild-type lines used were obtained from the Zebrafish International Resource Center (zebrafish.org).

### Cell culture and cell line generation

Cells were cultured at 5% CO2 and 21% O2. TC32 TC71 cells) were cultured using RPMI (Gibco) supplemented with 10% fetal bovine serum and 1% antibiotic antimycotic. NIH-3T3 cells were cultured using DMEM (Gibco) supplemented with 10% fetal bovine serum and 1% antibiotic antimycotic. TC32 shRNA lines cells were generated using the pZIP shRNA Promoter selection kit (transomics.com). EF1 shRNA targets the C-terminal region of Fli1 and the C shRNA is a scrambled sequence (Table S3). shRNA cells were cultured using RPMI (Gibco) supplemented with 10% fetal bovine serum and 1% antibiotic antimycotic. shRNA knockdown was induced by 1μg/ml doxycycline in cell culture 48 hrs prior to injection, and 10μg/ml doxycycline in E3 fish larval growth medium 24 hrs following injection, and shRNA knockdown was verified by western blotting using rabbit anti-Fli1 (AbCam ab15289) (Figure S3).

Fluorescent constructs were introduced into cells using the pLVX lentiviral system (Clontech) and selected using antibiotic resistance to either puromycin or neomycin. The GFP-FTractin and mRuby2-FTractin constructs contain residues 9–52 of the enzyme IPTKA30 fused to eGFP or mRuby2, respectively. pLenti6-H2B-mCherry was acquired from Addgene (generated by Wittmann lab, UCSF, Addgene plasmid #89766) [69].

### Xenografts

For xenografts of NIH-3T3, TC71, and TC32, preparation of cell mixture consisted of 99% or 90% mRuby2-Ftractin labeled cells and 1% or 10% GFP-FTractin labeled cells, for HBV and PVS, or CHT, respectively. These ratios were chosen to achieve optimum number of isolated cells not contacting other cells within a cluster. Cell mixtures (2×10^6^ cells) were run through a 70um cell strainer and resuspended in 5% FBS mixture for optimal cell viability.

Zebrafish embryos were collected, dechorionated and treated with 0.1 mM phenylthiourea (PTU) starting at 24 hpf to prevent pigmentation. Xenografts were generated by microinjection of cell mixture into zebrafish larvae (2 dpf) with glass capillary needles into PVS or HBV. For each cell xenograft group, cells were injected into 3-aminobenzoic acid ethyl ester (Tricaine)-anesthetized embryos. For PVS injections, larvae with cells disseminated to the CHT through passive pressure via the vasculature during injection were identified under a fluorescent stereomicroscope 1 hour post injections. This helped us (1) to increase the throughput of imaging by having two seeding sites within one embryo and (2) to select specifically for seeding-site specific cell adaptation rather than selection of a potentially more migratory population from the primary site of injection. Injected zebrafish larvae were incubated for 1-2 days in 34 °C before imaging.

### Immunofluorescence

For immunostaining of zebrafish larvae, anesthetized embryos were fixed in 4% paraformaldehyde/1XPBS for 2hrs in scintillation vials, rinsed in PBS +0.5% Tween (PBST), and stored in PBS up to two weeks. Immunostaining was carried out as previously described [70]. Primary antibodies were rabbit anti-GFP (AbCam, ab290) and mouse anti-Phospho-Histone H3 (Cell Signaling, 9706S). TUNEL assay was performed using the ApopTag Green In-Situ Apoptosis Detection Kit (Millipore). Xenografted fish were immunostained for GFP prior to high resolution imaging of single cells.

### Human Tumor Histology

Ewing Sarcoma tissue samples were obtained as part of standard clinical management. All histology samples were evaluated by an expert sarcoma pathologist.

### Imaging

Light-sheet fluorescence microscopes were used to image the zebrafish larvae. Fixed zebrafish larvae were embedded in 1% low-melting agarose into sonicated, ethanol-washed Fluorinated Ethylene propylene (FEP) tubes (Zeus, HS 029EXP/ .018 REC). The tube is held by a custom-made mount, which allows 3D translation and rotation in our light-sheet microscopes.

For high-resolution 3D imaging of cell morphologies, axially swept light-sheet microscopy (ASLM) was developed as previously described [46, 71]. Briefly, a 25X, NA1.1 objective water-dipping (CFI75 Apo 25XC W, Nikon Instruments) is used for detection, in combination with a 500 mm achromatic convex lens (49-396, Edmund Optics) forms the image with a total magnification of 62.5X on an sCMOS camera (ORCA-Flash4.0, Hamamatsu Photonics). A 28.6X NA0.7 water-dipping objective (54-10-7@488-910nm, Special Optics) is used for light-sheet illumination. In our ASLM set-up, the resolution is approximately 300 nm laterally and 450 nm axially over a 200×100 μm^2^ field of view (FOV).

For imaging of immunostained fish larvae, a conventional multidirectional selective plane imaging microscope (mSPIM) [72] was used as previously described [73]. Briefly, a 16X NA 0.8 water-dipping objective (CFI75 LWD 16XW, Nikon Instruments) is used for detection. In combination with a 200 mm tube lens (58-520, Edmund Optics) forms the image on an sCMOS camera (ORCA-Flash4.0, Hamamatsu Photonics). The lateral resolution is approximately 350-400 nm. A 10X NA 0.28 objective (M Plan Apo 10x, 378-803-3, Mitutoyo) is used for light-sheet illumination. The thickness and the length of the light-sheet are controlled by a variable slit (VA100C, Thorlabs) placed at the conjugate plane of the back focal plane of the illumination objective. Usually, we adjust the variable slit to create a light-sheet with a thickness of 3-4 μm, which has a length of approximately 400 μm to cover a large FOV. Under these conditions, the axial resolution is estimated at 1.4 μm.

### Image Processing

For high resolution 3D image processing, regions of interest around individual cells were detected automatically usingthresholding of the smoothed image (sigma=5, thresholded at 5%) and computing the connected components to isolate volumes of single cell image volumes from the raw data image. In some cases, additional user-aided cropping is done using ImageJ. We then applied drift correction using the StackReg plugin [74] in ImageJ [75] to counter sample movement in Z during acquisition, via MIJ, a java package for running ImageJ within MATLAB. Finally, we applied deconvolution to all images prior to segmentation, using blind deconvolution with a synthesized PSF with 10 iterations in MATLAB.

### Segmentation and Geometric feature extraction

For 3D morphology classification, we employed a 3D morphology pipeline described in Driscoll *et* al. [12] to segment cell volumes from deconvoluted single cell image volumes. For the segmentation process, we smoothed the deconvoluted images with a 3D Gaussian kernel (σ = 3 pixels) and applied a gamma correction (ϒ = 0.7), then employed an Otsu threshold, followed by a 3D grayscale flood-fill operation to fill possible holes. Disconnected parts from the main image were removed. The binary images of all cells were visually inspected for segmentation artifacts. Miss-segmented cells were excluded from the data set.

Computation 3D shape descriptors require parameters including volume, surface area, 3D convex hull, minimum circumscribed sphere and maximum inscribed sphere. The basic shape descriptors of segmented cells were computed using ‘regionprops3’ in MATLAB. Other features were calculated based on the formulas indicated in Table S1. These features include but are not limited to roughness, various definitions of sphericity, and 3D aspect ratio. The post-processing, segmentation, and geometric feature extraction pipelines are accessible via the Danuser lab GitHub [76].

For location within clusters, the region of interest detectors described above were reused to isolate segmented volumes. Individual cell surfaces were extracted from cell volumes using *Surfaces maps* from 3D Object Counter plugin in Fiji [77]. Measurements of percent of contact with other cells were made based on measurements of colocalization of the surfaces of the subsequent binary images. These were compared to individual geometric features as captured by the 3D morphology pipeline described above.

For quantification of proliferation and apoptosis within clusters, we utilized manually-aided threshold based segmentation followed by 3D Objects Counter plugin [77] in ImageJ. Size of cluster or touching objects was estimated by dividing the measured volume of the object by the measured volume of an isolated object (e.g. PH3-labeled nucleus) within the same fish.

For quantification of change in cluster size over three days (Figure S2a-d), segmentation of areas of interest and measurements of changes in area were done in Cell Profiler [78].

### Statistics and Visualizations

Violin plots, boxplots, and scatter plots of individual features at the cell-level or fish-level and corresponding statistical tests were performed in GraphPad Prism 9 (La Jolla, Ca). Outliers for individual parameters were found by ROUT method (Q = 1%) and excluded from the analysis. To address sample variability, fish level data points were generated based on the median of each geometrical feature per fish per region. Sample sizes are provided in figure legends.

For multivariable statistical analysis, Principal Component Analysis (PCA), scatter plots of PCs, and biplot were applied or generated in MATLAB. For visualization purpose, we defined the geometrical classes using the unsupervised learning k-means clustering algorithm in 2D PCA space and calculated the contribution of each geometrical class in each experimental condition by hierarchical clustering. Bagplots were generated in MATLAB as described in Verboven and Hubert, 2010 [79].

Similarity matrices were based on pairwise permutation p-values between different conditions. The metric used for permutation test (n=1500 iterations) was the Tukey median of each distribution, as defined by [48].

3D surface renderings of single cells were created with ChimeraX [80] as described in *Driscoll et al*. [12]. Rendering of microscopy set-up was created with AutoCad (Autodesk). 3D rendering in Figure S2 was created with Amira software (ThermoFisher). Schematics were created with BioRender.com.

## Acknowledgments

We would like to thank Meghan Driscoll, Erik Welf, David Saucier, Jungsik Noh, Jaewon Huh, and Marcel Mettlen for fruitful discussion and help with image analysis. We would also like to thank David McFadden (inducible shRNA lines), Ralph Deberardinis (TC32 cell line), Sandra Schmid (NIH-3T3 cell line), and Angelique Whitehurst (TC71 cell line) labs (UTSW) for sharing reagents. Funding for this work in the Danuser lab has been provided through R35 GM136428 (NIH). Dagan Segal acknowledges funding from RP160157 (CPRIT) and EMBO Long-term Research Fellowship (ALTF 626-2018). Reto Fiolka acknowledges funding by the NIH (grants R33 CA235254 and R35 GM133522). James Amatruda acknowledges funding from the 1 Million 4 Anna foundation.

## Author Contributions

Conceptualization, D.S., G.D, J.F.A, Methodology D.S., H.M., B.C., Investigation, D.S., H.S.,B.C.,P.R.,M.W., D.R. Writing, D.S., G.D, and J.F.A, Funding Acquisition D.S., G.D., J.F.A, and R.F., Resources, G.D., J.F.A, and R.F., Supervision, G.D., and J.F.A.

## Supplemental Information

**Figure S1:**
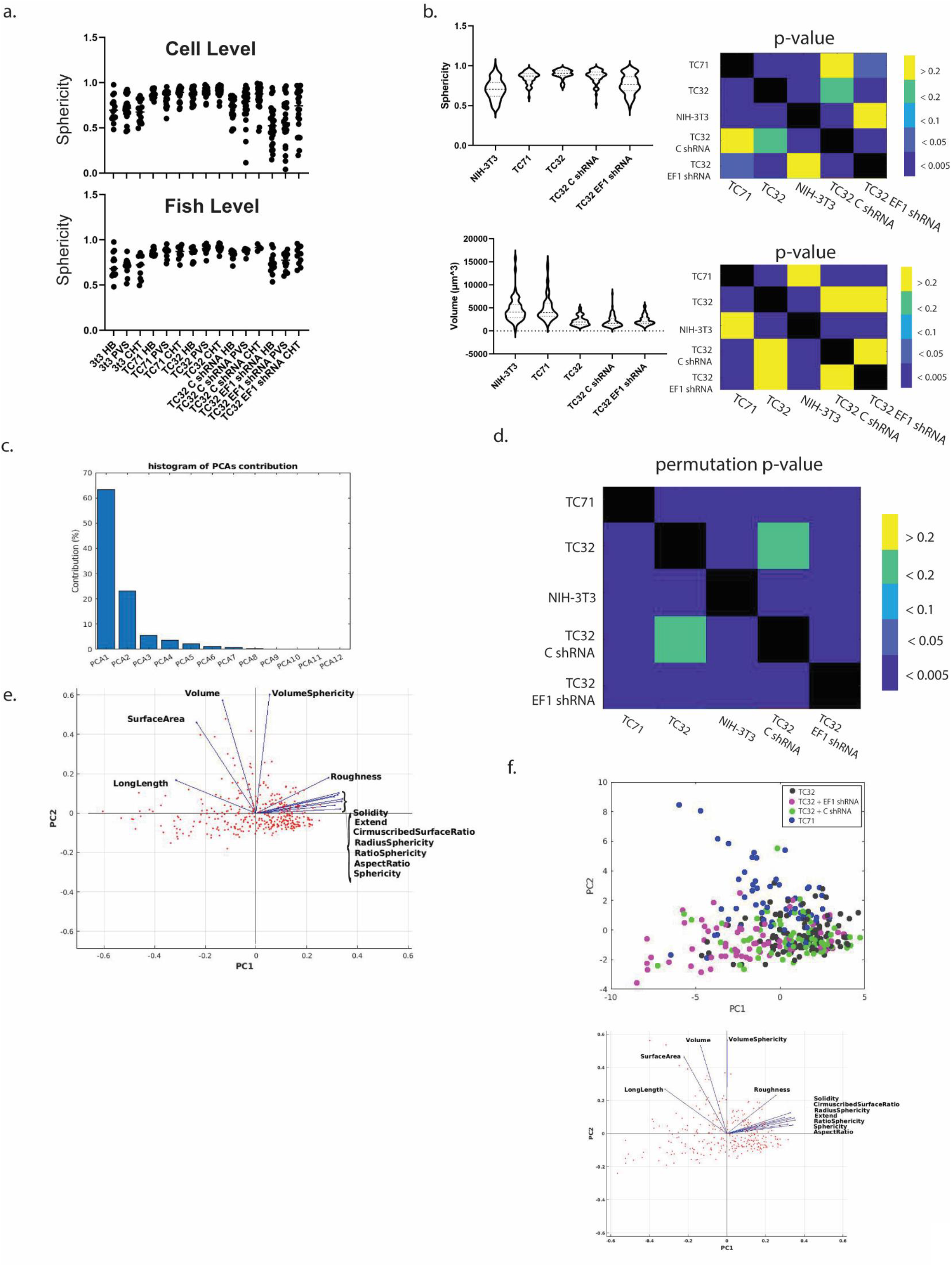
Classification of morphologies captures heterogeneity of cell states in EwS cells. **(a)** Sphericity single-cell level (n=15,22,15,19,23,18,19,41,37,18,25,28,25,26,19 for HBV/PVS/CHT of NIH-3T3, TC71, TC32, TC32 C shRNA, TC32 EF1 shRNA, respectively) or fish-level (n=13,11,10,12,11,12,11,16,13,10,7,4,14,12,10 for HBV/PVS/CHT of NIH-3T3, TC71, TC32, TC32 C shRNA, TC32 EF1 shRNA, respectively) show similar distributions, with low fish-to-fish variability. Significant shifts between bins already for a single feature at the fish level indicates that cell type, molecular condition, and seeding site play a role in the control of cell morphogenesis (ANOVA, p < 0.0001). **(b)** Violin plots (left) and similarity matrices displaying p-values (right, Kruskal-Wallis test, post-hoc Dunn’s test) of two prominent individual features, sphericity (top) and volume (bottom). Individual features alone cannot capture the full heterogeneity of the dataset (n = 52, 60, 97,71,70 for NIH-3T3, TC71, TC32, TC32 C shRNA, TC32 EF1 shRNA, respectively). **(c)** Histogram of the relative contributions of principal components (PCs). The first two PCs capture 86% of the heterogeneity in the data and are used for further analysis. **(d)** A similarity matrix based on pairwise permutation tests of the tukey median values of the first 2 PCs shows significantly different distributions between all experimental groups except TC32 and TC32 C shRNA. **(e)** A biplot displaying the relative contribution of each geometric feature to the first 2 PCs. All features appear to contribute to the first two PCs, with sphericity and volume strongly influencing PCs 1 and 2, respectively. **(f)** When excluding outlier group NIH-3T3, the PCA space (top) and contribution of features (bottom) remain largely the same while the classes become shifted, suggesting the PCA for the 12 3-dimensional global features defined constitutes a robust space for the definition of cell morphological classes.

**Figure S2:**
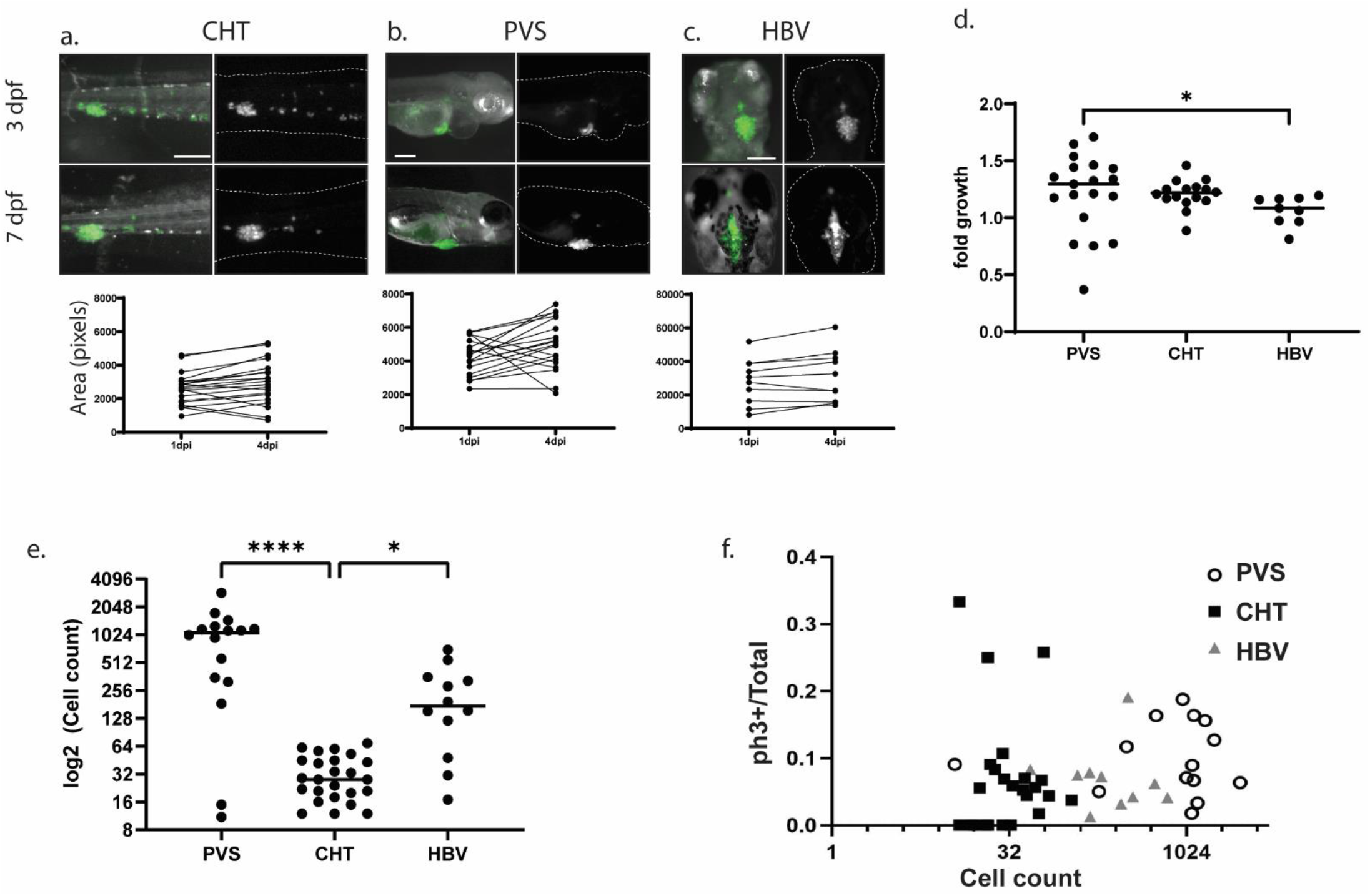
Proliferation but not apoptosis changes as a function of tissue microenvironment. **(a-c)** Images of TC71-GFP cells xenografted in zebrafish larvae at 2 days post fertilization (dpf) and imaged in their respective seeding sites (CHT, PVS, HBV) 3 and 7 dpf. Graphs indicate change in area over 4 days. Note that individual paired measurements within each tissue ME suggest that fold growth change is not influenced by initial seeding cluster size. **(d)** Fold growth as measured by change in measured area of CHT (a), PVS (b), or HBV (c) over a period of 4 days. PVS shows a larger increase in fold growth compared to HBV. **(e)** Relative size of TC71 clusters varies greatly as a function of tissue ME, which is likely a result of differential seeding rates during xenografts. **(f)** XY plot of cluster size (cell count) vs. proportion of PH3 positive cells (see figure 3). Proliferation rate does not seem to correlate to size of clusters, suggesting this difference does not greatly impact functional state of cells. Asterisks mark significant differences (Dunn’s multiple comparison test, p<0.05). Scale bars 20 um.

**Figure S3:**
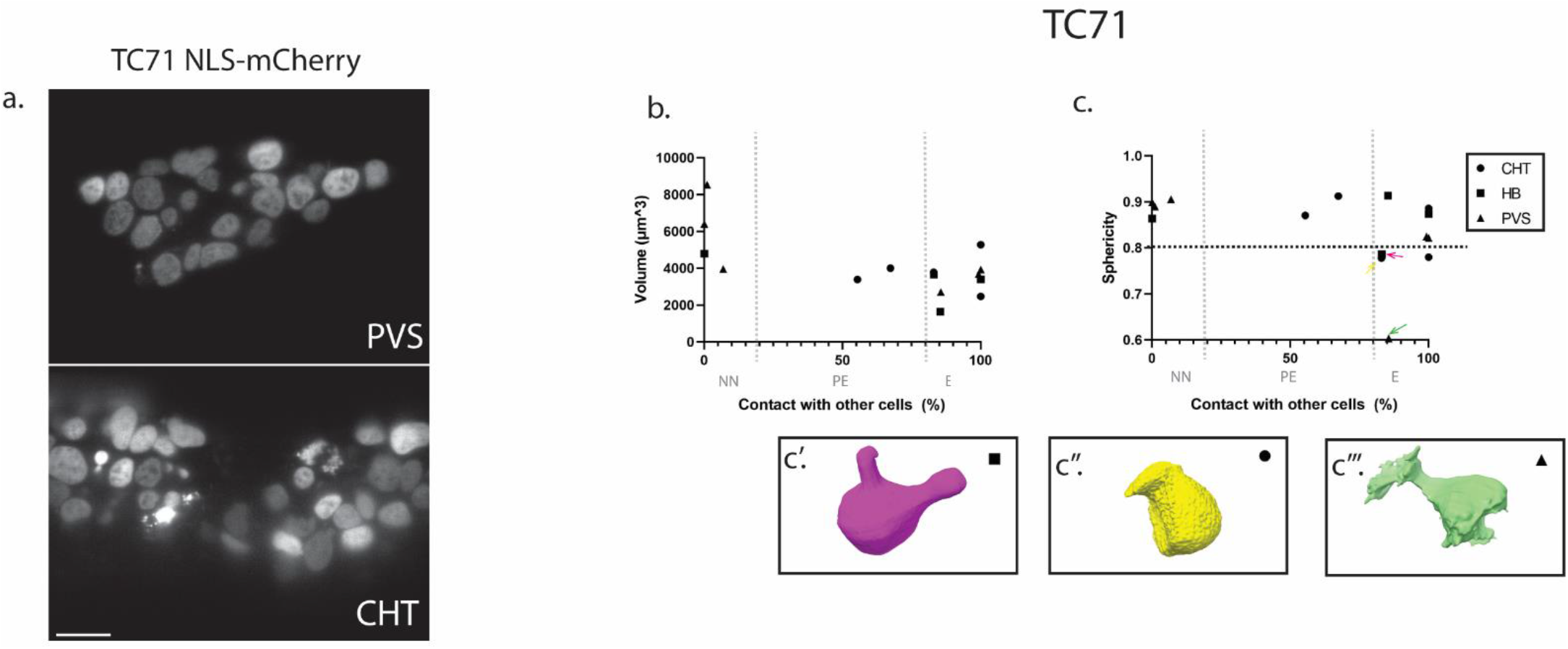
Tissue architecture affects cell state. **(a)** No pseudorosette-like structures are typically observed in single optical slice of TC71 cells expressing nuclear marker H2B-Cherry in the perivitelline space (PVS) or caudal hematopoetic tissue (CHT), as opposed to the HBV (see figure 5e). **(b-c)** Proportion of contact with other TC71 cells is plotted against Volume (b) and Sphericity (c). While no clear relationship is observed for volume, a few *embedded* (E) cells but not other groups show cells with distinctly low sphericity (>0.8). Arrows indicate cells from all three regions with distinctive cell morphological states, rendered in c’-c’’’ and color-coded by k-means clustering in Figure 2e. Scale bars 20um.

**Figure S4:**
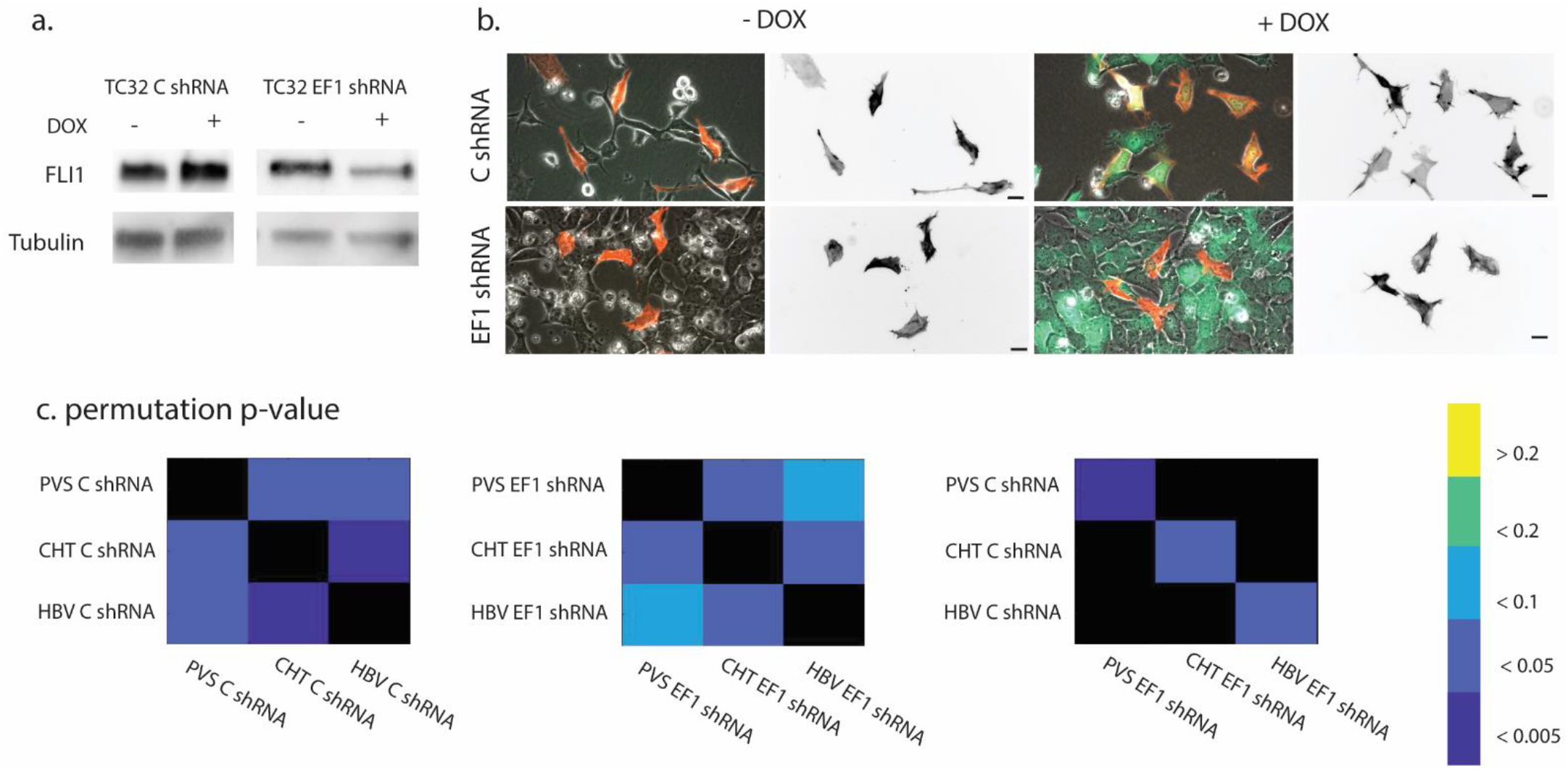
Knockdown of EF1 causes variable changes in morphologies as a function of tissue microenvironment. **(a)** Western blot shows efficient knockdown of EWS-FLI1 in cell culture conditions upon 72hrs addition of doxycycline (DOX). **(b)** Images of TC32 cells expressing C shRNA or EF1 shRNA with or without addition of DOX in cell culture. A subset of cells express F-Tractin mRuby2 (red on left, grey on right), while all cells express the DOX-inducible cytoplasmic GFP, allowing for mosaic cell labeling. Mild changes in actin organization are observed with expression of EF1 shRNA. However, no clear distinctions in cell morphology are seen. **(c)** Similarity matrix based on pairwise permutation test of Tukey depth median of first two PCs shows distinct differences in cell states between all three tissue MEs in TC32 C shRNA (left). Upon loss of EF1, cell states in all three regions change significantly (right), and enviromentally-dependent differences become blurred and less distinct (middle), suggesting expression of EF1 enables tissue-specific specialization of cells.

**Figure S5:**
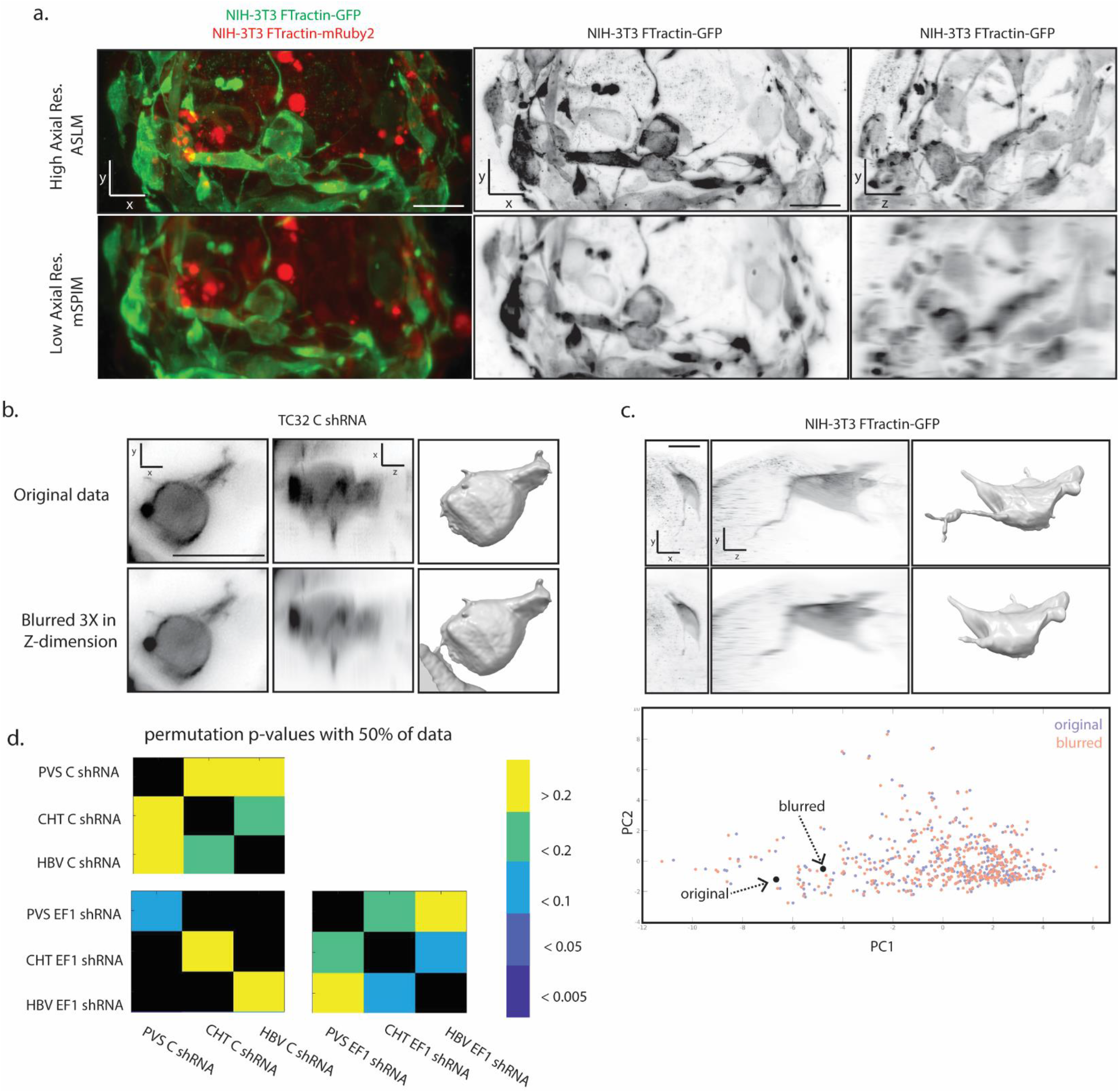
Improved imaging and analytical techniques are required for quantitative analysis of cancer cells *in situ*. **(a)** The axially swept light sheet microscope (ASLM) increases contrast and significantly improves axial resolution when compared to the multidirectional selective plane illumination microscopes (mSPIM) often utilized in imaging of zebrafish. Shown are mixtures of flourescently labeled NIH-3T3 expressing FTractin-mRuby2 (90% of cells, red) or FTractin-GFP (10% of cells, green and grey) in the zebrafish hindbrain ventricle (HBV), 3 days post fertilization. Left to right: XY, XY, and ZY maximum intensity projections of images acquired on ASLM (upper panel) or mSPIM (lower panel). **(b-c)** Simulations of blurred data in Z-dimension causing non-isotropic resolution leads to segmentation artifacts, such as perceived contact with another cell (b) or loss of some protrusions (c, upper). While cells as depicted in b would be identified in quality control and removed from analysis, loss of protrusions such as seen in c would not be identified and lead to perceived shifts in PC space and loss of distinction between experimental groups (c, lower). **(d)** Loss of 50% of data leads to loss of distinction between experimental groups, as visualized by similarity matrix based on pairwise permutation tests (compare to Figure S4c).

**Movie S1: EwS cells form rosette-like structures in Hindbrain.** TC32 and TC71 cells expressing Ftractin-GFP (grey, green) or F-Tractin-mRuby2 (red), in zebrafish larval hindbrain ventricle (3dpf). In both EwS cell lines, 3D rotation of image z-stack shows pseudorosette-like polarization of cells, with cellular protrusions pointing towards a center of a sphere and cell bodies organized outwards. Running through the z-dimension of the optical slices illustrates how rosettes form within a subset of the clusters, rather than encompass its entirety.

**Movie S2: Rosette-like organization maintained upon loss of EF1.** TC32 cell expressing C shRNA or EF1 shRNA in zebrafish larval hindbrain ventricle (3dpf). A subset of cells express F-Tractin mRuby2 (grey, red), while all cells express the DOX-inducible cytoplasmic GFP (green). Rosette-like formation is sustained in the EF1 knockdown condition, despite gross morphological changes. See also Figure 4.

### Tables

**Table S1:** Table describing the mathematical definition of the 12 3-dimensional global shape features used for morphology classification.

| **Global geometrical features** | **Definition** |
| --- | --- |
| Volume (V) |  |
| Surface area (S) |  |
| Solidity | $V_{\text{Cell}} / V_{\text{Convex hull}}$ |
| Sphericity | $\pi^{1/3} (6V_{\text{Cell}})^{2/3} / S_{\text{Cell}}$ |
| Long length | Longest length in all direction |
| Extend | $V_{\text{Cell}} / V_{\text{Bounding box}}$ |
| Compactness | $S_{\text{Cell}}^{1.5} / V_{\text{Cell}}$ |
| Equivalent radius | Radius of a sphere with the same cell volume |
| Aspect ratio | $L_{\text{Short}} / L_{\text{Long}}$ |
| Roughness | $S_{\text{Cell}} / S_{\text{Convex hull}}$ |
| Volume sphericity | $V_{\text{Cell}} / V_{\text{Circumscribed sphere}}$ |
| Radius sphericity | Equivalent radius / $R_{\text{Circumscribed sphere}}$ |
| Spherical radius ratio | $R_{\text{Inscribed sphere}} / R_{\text{Circumscribed sphere}}$ |
| Circumscribed sphere area ratio | $S_{\text{Cell}} / S_{\text{Circumscribed sphere}}$ |

**Table S2:** Analytical framework is robust to differences in segmentation parameters. Table showing pair wise permutation p-values based on Tukey median (n = 1500 iterations) across parameter values. Note stability of p-values within 30% variation in parameter values, except in few cases when signal-to-noise ratios were high (highlighted in yellow), leading to sensitivity to underfiltering. Note also changes in number of processed cells due to variation in segmentation errors. Gray column indicates parameters used for analysis, which remained constant for all cells analyzed.

| | $\sigma = 2 \gamma = 0.70$ | $\sigma = 4 \gamma = 0.70$ | $\sigma = 3 \gamma = 0.55$ | $\sigma = 3 \gamma = 0.85$ | $\sigma = 3 \gamma = 0.70$ |
| --- | --- | --- | --- | --- | --- |
| 'CHT4300-PVS4300' | 0.091 | 0.009 | 0.123 | 0.065 | 0.017 |
| 'HB4300-PVS4300' | 0.000 | 0.019 | 0.002 | 0.013 | 0.011 |
| 'CHT4300-HB4300' | 0.000 | 0.000 | 0.000 | 0.000 | 0.000 |
| 'CHT 226-PVS 226' | 0.086 | 0.069 | 0.035 | 0.025 | 0.033 |
| 'CHT 226-HB 226' | 0.003 | 0.021 | 0.003 | 0.002 | 0.005 |
| 'PVS 226-HB 226' | 0.026 | 0.105 | 0.142 | 0.091 | 0.077 |
| 'CHT 32-PVS 32' | 0.585 | 0.271 | 0.619 | 0.714 | 0.570 |
| 'CHT 32-HB 32' | 0.001 | 0.000 | 0.000 | 0.001 | 0.001 |
| 'HB 32-PVS 32' | 0.008 | 0.004 | 0.003 | 0.003 | 0.002 |
| 'CHT 71-PVS 71' | 0.017 | 0.007 | 0.023 | 0.033 | 0.018 |
| 'CHT 71-HB 71' | 0.033 | 0.095 | 0.127 | 0.173 | 0.094 |
| 'HB 71-PVS 71' | 0.015 | 0.204 | 0.121 | 0.198 | 0.161 |
| 'CHT 33-PVS 33' | 0.611 | 0.524 | 0.398 | 0.370 | 0.356 |
| 'CHT 33-HB 33' | 0.902 | 0.927 | 0.849 | 0.891 | 0.755 |
| 'HB 33-PVS 33' | 0.637 | 0.523 | 0.648 | 0.494 | 0.673 |
| '33-32' | 0.000 | 0.000 | 0.000 | 0.000 | 0.000 |
| '33-71' | 0.000 | 0.000 | 0.000 | 0.000 | 0.000 |
| '33-226' | 0.000 | 0.000 | 0.000 | 0.000 | 0.000 |
| '33-4300' | 0.000 | 0.000 | 0.000 | 0.000 | 0.000 |
| '32-71' | 0.000 | 0.000 | 0.000 | 0.000 | 0.000 |
| '32-226' | 0.000 | 0.000 | 0.000 | 0.000 | 0.000 |
| '32-4300' | 0.699 | 0.429 | 0.603 | 0.549 | 0.471 |
| '4300-226' | 0.000 | 0.000 | 0.000 | 0.000 | 0.000 |
| '4300-71' | 0.000 | 0.000 | 0.000 | 0.000 | 0.000 |
| '71-226' | 0.000 | 0.000 | 0.000 | 0.000 | 0.000 |
| 'CHT4300-CHT 226' | 0.009 | 0.002 | 0.051 | 0.007 | 0.014 |
| 'HB4300-HB 226' | 0.016 | 0.017 | 0.036 | 0.028 | 0.029 |
| 'PVS4300-PVS 226' | 0.006 | 0.001 | 0.001 | 0.001 | 0.003 |
| **Number of Cells Analyzed** | **341** | **338** | **333** | **348** | **348** |

**Table S3:**
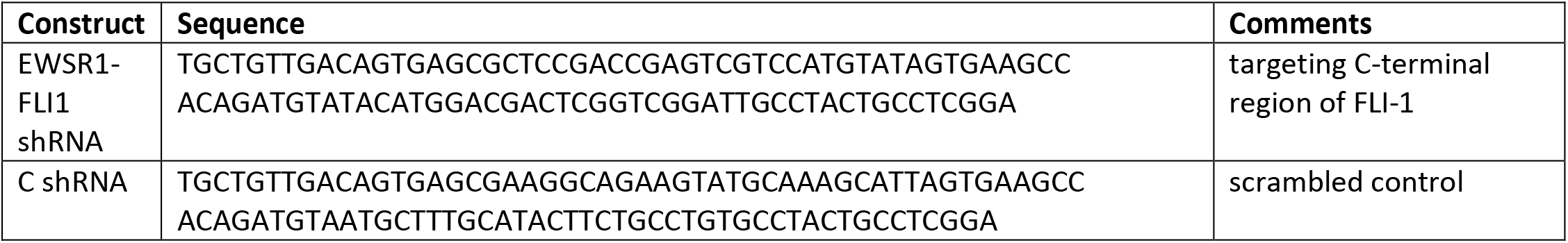
Sequences for EWSR1-FLI1 and scrambled-targeting shRNA constructs.

